# Free-living Bacterial Communities Are Mostly Dominated by Oligotrophs

**DOI:** 10.1101/350348

**Authors:** Yingnan Gao, Martin Wu

## Abstract

In response to resource availability, bacteria have evolved two distinct ecological strategies. Copiotrophic bacteria grow fast and are heavily favored by selection where the resource is abundant. In contrast, oligotrophic bacteria grow slowly but more efficiently and are highly adaptive in nutrient-poor environments (Koch, 2001). Although oligotrophs and copiotrophs are ubiquitous, except for a few well-characterized environments like the open ocean and animal gut, the relative abundance of oligotrophic and copiotrophic bacteria and their importance in the global ecosystem are still unclear. In addition, although several studies have demonstrated the impact of nutrient availability on the bacterial community structure under experimental conditions (Klappenbach et al., 2000, Nemergut et al., 2016), the role of nutrients in shaping the structures of bacterial communities in their natural habitats remains largely unknown. Using the ribosomal RNA operon (*rrn*) copy number to capture the bacterial ecological strategy, we analyzed 44,045 samples from two large bacterial community repositories that cover 78 environmental types. Here we show that animal-associated microbiota are dominated by copiotrophs while plant-associated and free-living bacterial communities are mostly dominated by oligotrophs. Our results suggest that nutrient availability plays an important role in determining the structure and ecological strategy of bacterial communities in nature. We demonstrate that the average and distribution of *rrn* copy number are simple yet robust predictors of the ecological strategy of bacterial communities that can be applied to all sequence-based microbial surveys to link the community structure and function.

## Introduction

The fundamental trade-off between the growth rate and efficiency in bacteria leads to genomic features that distinguish between copiotrophs and oligotrophs (Lauro et al., 2009; Roller et al., 2016). These features include genome size (Roller et al., 2016), codon usage bias (Vieira-Silva & Rocha, 2010), transporter gene diversity (Lauro et al., 2009), motility (Lauro et al., 2009; Roller et al., 2016) and etc. Among them, the copy number of ribosomal RNA operon (*rrn*) is the best studied genomic trait that predicts the ecological strategy of bacterial species. The *rrn* copy number positively correlates with cellular ribosomal content and the maximum growth rate (Roller et al., 2016; Vieira-Silva & Rocha, 2010), and deletion of *rrn* reduces the maximum growth rate of *Escherichia coli* and *Bacillus subtilis* (Nanamiya et al., 2010; Stevenson & Schmidt, 2004; Yano et al., 2013). On the other hand, the *rrn* copy number negatively correlates with the growth efficiency. One study has shown that the carbon use efficiency and protein yield (protein synthesized per unit O_2_ consumed) decreases when the *rrn* copy number increases in several strains of bacteria (Roller et al., 2016). As such, high *rrn* copy number indicates a copiotrophic strategy while low *rrn* copy number predicts an oligotrophic one.

As bacteria adopt different ecological strategies, the *rrn* copy number in their genomes varies from 1 to as many as 15 (Klappenbach et al., 2000). Bacteria have the potential to rapidly change their *rrn* copy numbers in adaptation to the growth condition, as mutant strains of *B. subtilis* with only one copy of *rrn* increased their *rrn* copies through operon duplication within generations (Yano et al., 2013). Despite the potential for rapid change, there is clearly phylogenetic signal in bacterial *rrn* copy number, as closely related species tend to have similar *rrn* copy numbers (Kembel et al., 2012). The existence of such phylogenetic signal suggests that *rrn* copy number, albeit labile, is under strong natural selection. Although *rrn* copy number cannot be directly measured for environmental bacteria, the phylogenetic signal in *rrn* copy number makes it possible to estimate the copy number from the 16S rRNA gene sequence alone (Kembel et al., 2012). And such method is applicable to both 16S rRNA amplicon and shotgun metagenomic sequence data, making prediction of ecological strategies of individual bacterial species and the whole community relatively simple and straightforward.

## Results

First we set out to identify an optimal *rrn* copy number that can be used to classify oligotrophs and copiotrophs. We used 63 representatives of oligotrophs and copiotrophs classified by Lauro *et al* (Lauro et al., 2009) as our training set. As expected, oligotrophs and copiotrophs are well separated in their *rrn* copy number distribution (Figure 1A). The area under the curve (AUC) of the receiver operating characteristic (ROC) curve is 0.86 (Figure 1B), indicating that classification by *rrn* copy number has high accuracy (Šimundić, 2008). Maximizing Youden’s J statistic, we found that 2 was the optimal cutoff to distinguish oligotrophs and copiotrophs, with a true positive rate of 0.789 and a false positive rate 0.200 for identifying oligotroph. Using this cutoff, we classified bacterial species (Operational Taxonomic Units or OTUs) as oligotrophs if they had 1 or 2 *rrn* copies or as copiotrophs if they had 3 or more *rrn* copies.

**Figure 1.**
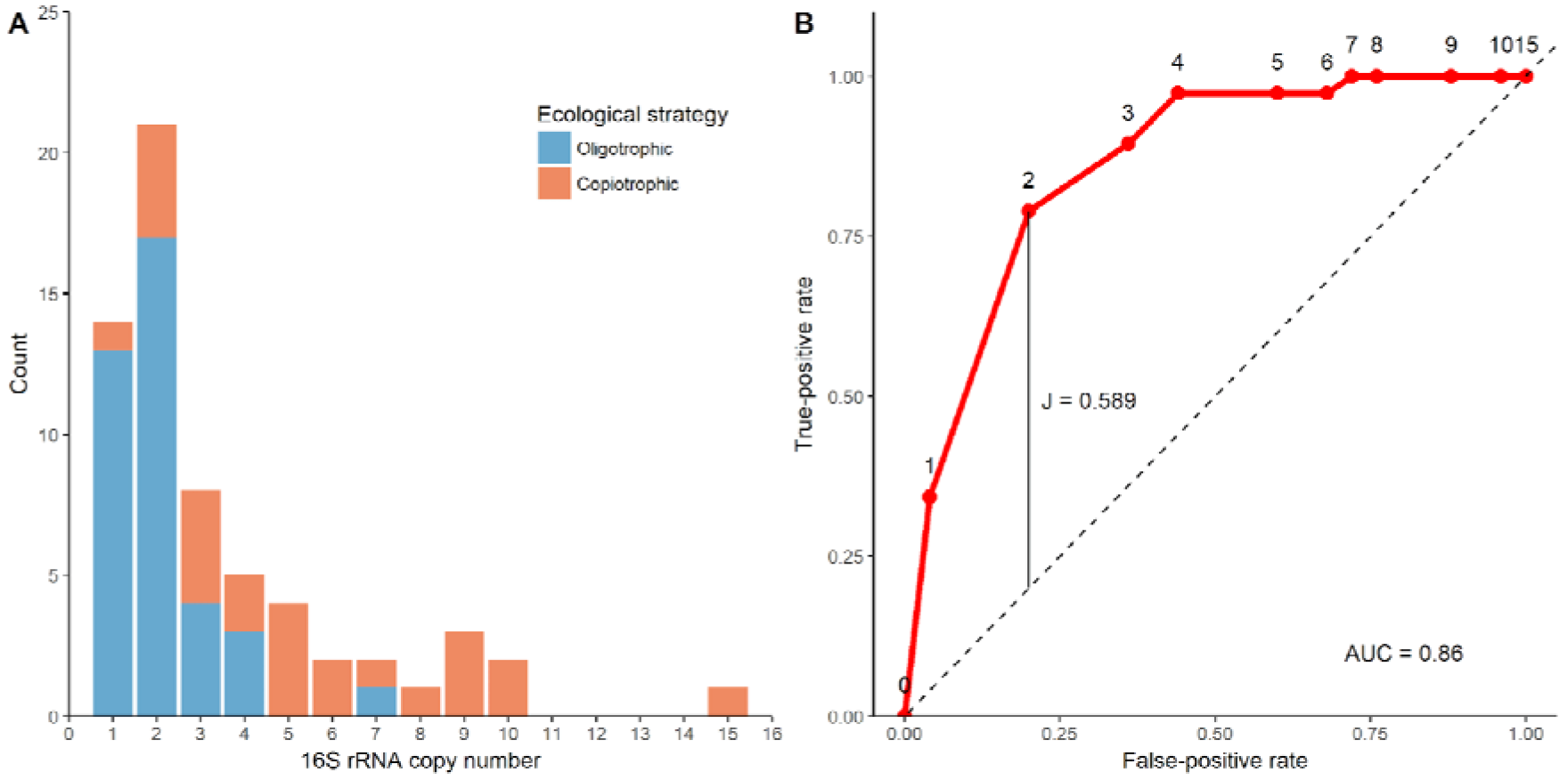
(A) The distribution of *rrn* copy number of the 63 representative species that were identified as either oligotrophic (blue) or copiotrophic (orange). (B) The ROC curve for using the *rrn* copy number to distinguish oligotrophic and copiotrophic bacteria, with an AUC of 0.86. The best copy number cutoff is 2 with a maximum Youden’s J statistic of 0.589.

The free-living bacterial community in open ocean is known to be dominated by oligotrophic bacteria (Lauro et al., 2009; Yooseph et al., 2010). Therefore, we tested whether the average copy number (ACN) and distribution of *rrn* captured the ecological strategy of marine bacteria in the open ocean using the shotgun metagenomic sequencing data from the Tara Oceans Expedition (TARA) (Pesant et al., 2015). The ACN in surface water samples from TARA ranged from 1.3 to 1.9, with a median of 1.4 (Supplementary Figure 1A). As expected, the community is dominated by oligotrophic bacteria with 1 or 2 *rrn* copies. Overall, 93% of the bacterial cells in the community were oligotrophic, with 66% of the bacteria cells had only one copy of *rrn* and 27% of them had 2 copies (Supplementary Figure 1B).

Next we examined the ACN of bacterial communities from a broader range of environments present in the EBI Metagenomics dataset (Mitchell et al., 2018). In total, we analyzed 2,528 whole genome shotgun (WGS) metagenomic samples that covered 6 environmental categories (freshwater, marine, non-marine saline, soil, animal and human-associated) and 21 environmental types (Supplementary Table 1). We observed that ACN varied substantially between environmental categories (Figure 2A). Animal and human-associated bacterial communities had the highest ACN (median 4.4 and 4.8 respectively), while those from marine, non-marine saline and soil environments had the lowest ACN (median 1.8, 2.0 and 2.0, respectively). Freshwater bacteria communities had intermediate ACN (median 3.1).

**Figure 2.**
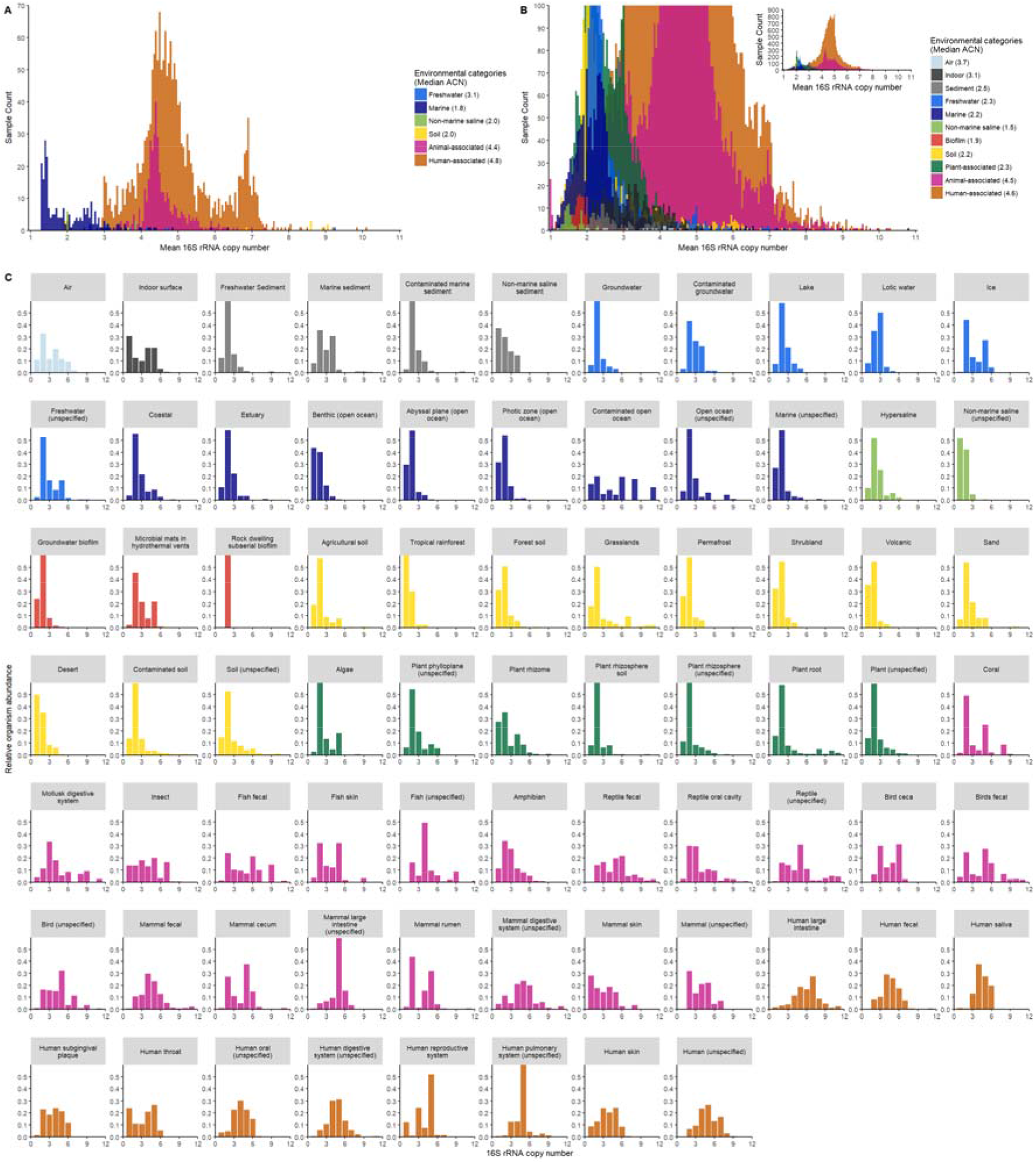
Distribution of the ACN of 2,528 samples surveyed by WGS metagenomics (A) and 41,517 samples surveyed by 16S rRNA amplicon sequencing (B) in the EBI Metagenomics and the EMP databases. The relative abundance of bacterial species was plotted against their *rrn* copy number for the amplicon sequencing samples (C). Samples were colored by the environment categories.

As the majority of microbial surveys are based on 16S rRNA amplicon sequencing, we investigated whether the ACNs estimated from amplicon sequencing were consistent with those estimated from WGS sequence data, out of concern that PCR bias associated with amplicon sequencing (Acinas et al., 2005) can potentially skew the ACN. From the EBI Metagenomics database, we identified 275 animal and human-associated microbiomes that were surveyed by both methods. We compared the ACNs estimated from the WGS and amplicon sequences of these 275 samples and found that they were highly correlated (Figure 3). The r-squared of the one-to-one fit between the two ACNs was 0.708 and the difference in ACNs estimated by the two methods was negligible (1.2% difference on average).

**Figure 3.**
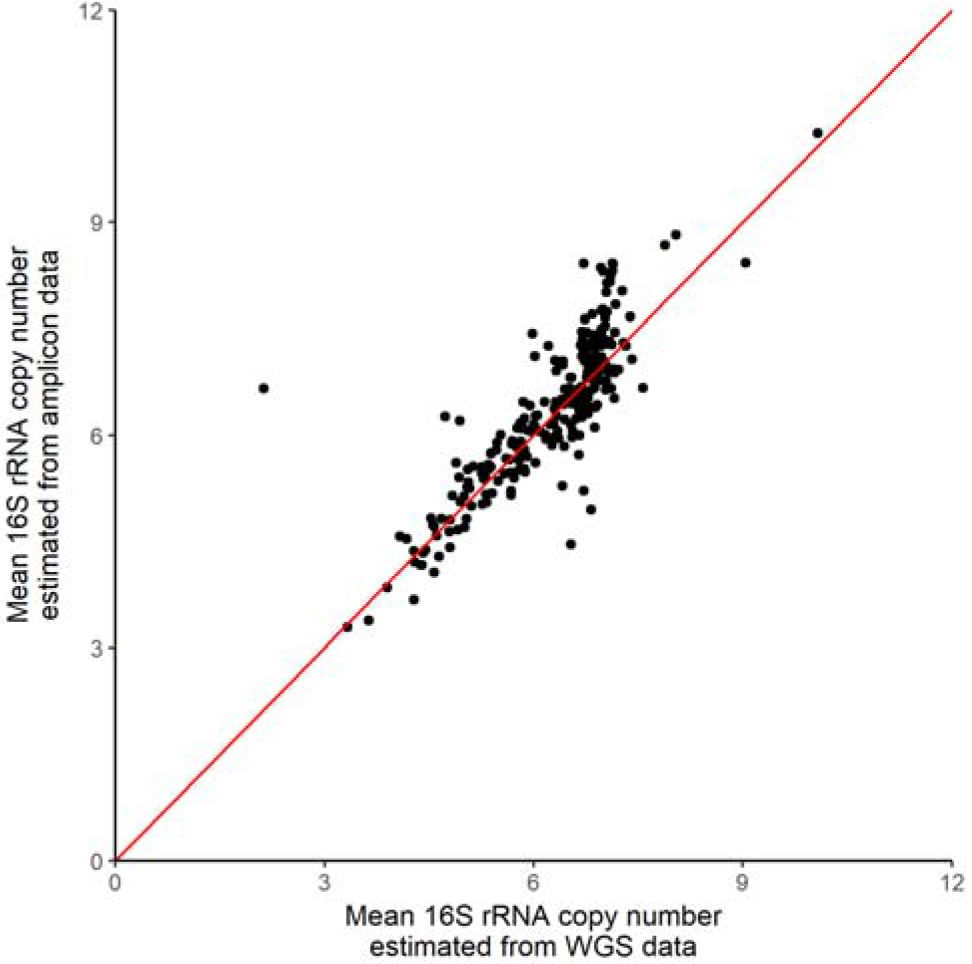
The ACN of 275 animal and human-associated bacterial communities surveyed by both 16S rRNA amplicon and WGS metagenomic sequencing methods. Red line illustrates the one-to-one fit between the ACN estimated from the two methods. The r-squared of the fit is 0.708.

Because ACN estimated from 16S rRNA amplicon data was reliable, we examined the ACN of bacterial communities surveyed by amplicon sequencing in both the EBI Metagenomics and Earth Microbiome Project (EMP) database (Thompson et al., 2017). In total, we analyzed 41,517 amplicon-sequenced samples from the two databases, which were 16 times larger than the WGS metagenomic data and together covered 11 environmental categories and 74 environmental types (Supplementary Table 1). ACN from the amplicon data varied between environment categories and showed a distribution pattern similar to what had been observed in the EBI WGS metagenomic data (Figure 2B) and a previous report (Thompson et al., 2017). Animal and human-associated bacterial communities had highest ACN (median 4.5 and 4.6 respectively), followed by air and indoor surface communities with intermediate ACN (median 3.7 and 3.1 respectively). Bacterial communities in the other environments (freshwater, marine, non-marine saline, soil, sediment, biofilm and plant-associated) all had low ACN (median less than 2.5).

We then examined the distribution of the *rrn* copy number within each of the 74 environmental types to directly assess the relative abundance of oligotrophs and copiotrophs in each type of environment. Figure 2C shows that animal and human-associated bacterial communities are dominated by copiotrophs (with 3 or more *rrn* copies) while free-living bacterial communities are dominated by oligotrophs (with 1 or 2 *rrn* copies). Plant-associated bacterial communities show a distribution of *rrn* copy number similar to those of the free-living ones, dominated by oligotrophs. We classified bacterial communities in which the relative abundance of oligotrophs was greater than 60% as oligotrophic, and communities in which the relative abundance of copiotrophs was greater than 60% as copiotrophic, and the rest of communities as intermediate. We found that 91.5% of 31,835 animal and human-associated bacterial communities were copiotrophic, while 72.6% of 6,215 free-living and 71.2% of 3,467 plant-associated microbiomes were oligotrophic (Figure 4).

**Figure 4.**
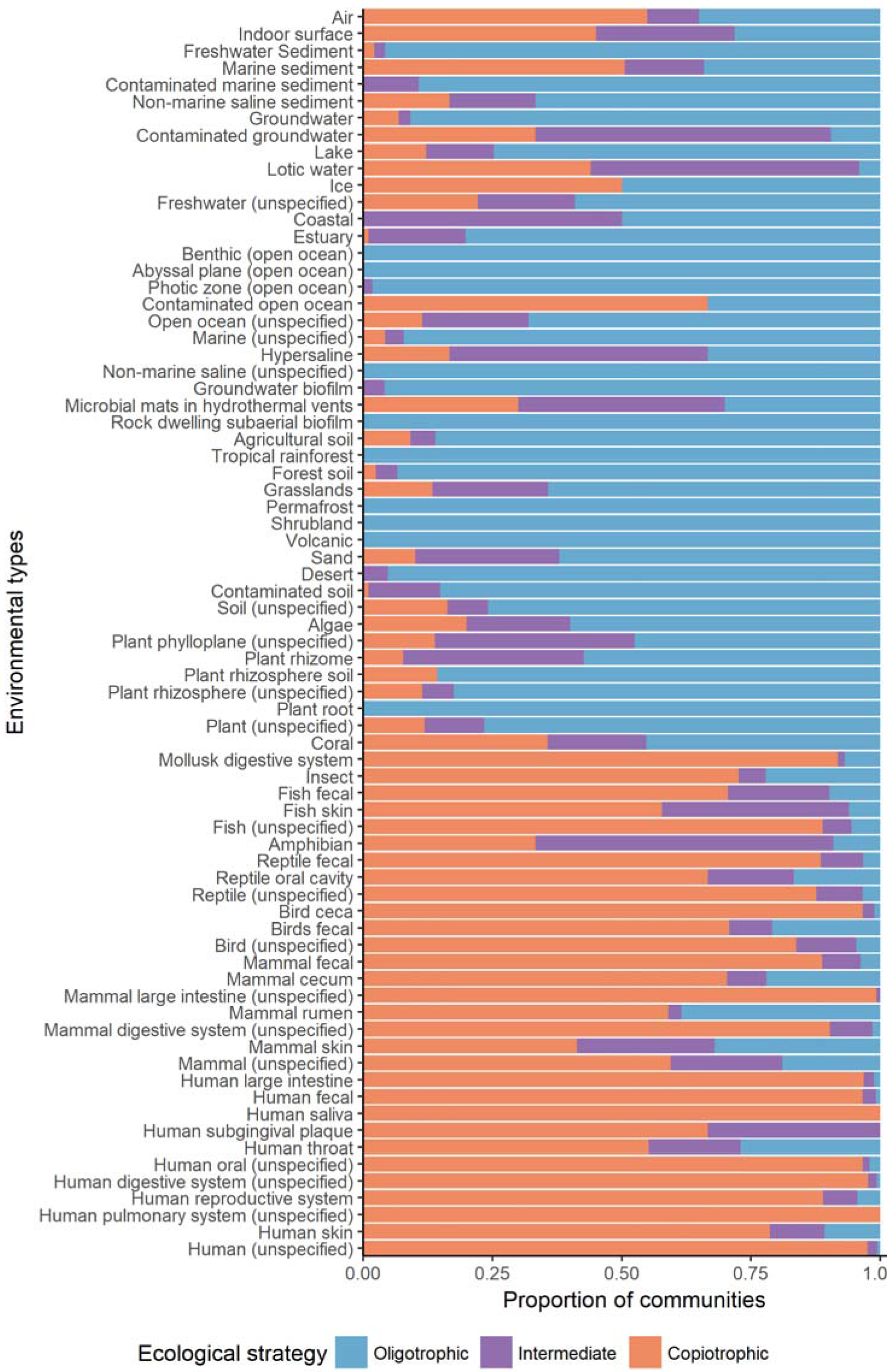
The proportion of 41,517 16S rRNA amplicon sequenced samples from 74 environmental types that are dominated (more than 60%) by oligotrophs (species with 1 or 2 copies or *rrn*, orange) or by copiotrophs (species with 3 or more copies of *rrn*, blue), or are intermediate (neither group was dominant, purple).

## Discussion

The *rrn* copy number has been shown to be a useful predictor of the ecological strategy of individual bacterial species (Klappenbach et al., 2000; Lauro et al., 2009; Roller et al., 2016) and bacterial communities (Nemergut et al., 2016; Thompson et al., 2017). Here we extended this line of research and used the distribution of *rrn* copy to reveal the existence of three types of bacterial communities in nature: those dominated by oligotrophs, copiotrophs or neither. Our analysis of a comprehensive collection of environmental samples indicates that most free-living and plant-associated bacterial communities are dominated by oligotrophs while animal and human-associated communities are dominated by copiotrophs. This pattern is consistent with our understanding of nutrient availability in the environment. In general, soil and water are considered nutrient poor (Barber & Lynch, 1977; Kuznetsov et al., 1979; Maeda et al., 2000; Ohta & Hattori, 1983) while animal-associated sites (gut, skin, etc) are considered nutrient rich. Despite the vast compositional differences among the communities analyzed in this study, the emergence of a global pattern in the relative abundance of oligotroph and copiotroph suggests that nutrient plays a key role in shaping the ecological strategy and the structure of the bacterial communities in nature. More generally, our results suggest that in nature selection exerts strong influence on the assembly of bacterial community compared to the other processes such as drift and dispersal. Interestingly, although the vast majority of groundwater samples were dominated by oligotrophs (90.9% of samples), only 9.5% of contaminated groundwater samples were dominated by oligotrophs. Similarly, the relative abundance of oligotrophs also decreased in the contaminated open ocean and soil. In addition, we did observe outliers in some environments. For example, Antarctic soil supplemented with organic carbon (Van Horn et al., 2014) had extremely high ACN (8.9) compared to untreated soil (3.1). Together, they provide compelling evidence that nutrient dictates the ecological strategy and the structure of the bacterial communities.

We have demonstrated that the average and distribution of *rrn* copy number are simple yet robust predictors of the ecological strategy of bacterial communities. Unlike other imprinted genomic features (Lauro et al., 2009) that require shotgun metagenomic sequences, *rrn* copy number can be estimated from 16S rRNA gene sequences alone and therefore is applicable to both shotgun metagenomic and 16S rRNA amplicon sequence data. As such, the metrics of average and distribution of *rrn* copy number can be applied in all sequence-based microbial surveys. Additionally, as the average copy number of *rrn* is weighted by the relative cell abundance, it will capture the overall ecological strategy of a community even at relatively low sequencing depth. These features make the community *rrn* copy number an extremely useful quantitative trait for studying the microbial ecosystem. By providing a link between the community structure and function, it adds values to 16S rRNA sequence data beyond simply quantifying species composition, and can help researchers generate hypotheses on how communities assemble in response to nutrient availability and other environmental factors. In addition, it can be used as proxies of the average growth rate and carbon use efficiency of bacterial communities to improve the current global carbon cycling models that have fixed values for these parameters, as has been suggested previously (Roller et al., 2016).

## Methods

### Classifying oligotrophic and copiotrophic bacteria with the *rrn* copy number

Using 43 genomic features, Lauro *et al* classified 126 strains of bacteria as either oligotrophic or copiotrophic (Lauro et al., 2009). We used these strains as a training set to identify an optimal *rrn* copy number cutoff for oligotroph/copiotroph classification. To reduce overrepresentation of certain clades, we selected one representative species from each of the 63 genera represented by the 126 strains and whose complete genome sequence was available (Supplementary Table 2). To evaluate the performance of using *rrn* copy number for oligotroph/copiotroph classification, we calculated the true-positive rate and the false-positive rate using each integer from 1 to 15 as the copy number cutoff, plotted the ROC curve and calculated the AUC. To find the optimal cutoff, we computed Youden’s J statistic for each cutoff, and selected the copy number that had the maximum Youden’s J statistic as the optimal cutoff. We then classified bacterial species whose *rrn* copy number was greater than the cutoff as copiotroph, and oligotroph otherwise.

### Compilation of Data

We used data from the Tara Oceans Expedition (Pesant et al., 2015) to test if the *rrn* copy number captured the oligotrophic ecological strategy of marine bacteria in the open ocean. The TARA dataset includes 139 bacterial community profiles derived from WGS metagenomic sequencing and focused specifically on the free-living bacterial communities in the open ocean across the globe. We downloaded the OTU abundance table and the environmental metadata from the TARA companion website (Sunagawa et al., 2015). We also downloaded the SILVA 16S rRNA reference sequences that were used in the TARA study to pick OTU at 97% similarity, and estimated their *rrn* copy numbers using the method described below. We analyzed 63 samples from the surface layer.

To explore the global ecological strategies of bacterial communities in nature, we compiled bacterial diversity survey data from two large microbial survey repositories, the EBI Metagenomics (Mitchell et al., 2018) and the EMP (Thompson et al., 2017). For the EBI Metagenomics dataset, we included 12,486 WGS metagenomic and 69,385 amplicon sequencing runs processed by the version 2.0 or 3.0 pipeline of the EBI Metagenomics (Mitchell et al., 2018). Sequencing runs of the same type (16S rRNA amplicon or WGS metagenomics) that were associated with the same sample were combined together. For the EMP dataset, we included 23,228 amplicon sequencing samples from its first release. We downloaded OTU abundance table and the metadata associated with each sample. The GreenGene 13.8 16S rRNA reference sequences were used for OTU picking at 97% similarity in both repositories. The *rrn* copy numbers of GreenGene reference sequences were estimated as described below. We filtered out WGS metagenomic runs whose 16S rRNA reads exceeded 5% of the total sequence reads and amplicon runs if less than 95% of the sequence reads were 16S rRNA reads, as these samples were likely 16S rRNA amplicon sequencing data mislabeled as WGS metagenomic data or *vice versa.* We then removed samples if less than 80% of their 16S rRNA reads could be mapped to the GreenGene reference sequences or if the total mapped reads were fewer than 400. We removed samples that only surveyed or were enriched for a specific bacterial clade. We also removed samples whose environmental types were too general or mislabelled, and environmental types that contained less than 5 samples. At the end, a total of 44,045 samples (2,528 WGS metagenomic and 41,517 amplicon sequencing) remained after the quality filtering. Together, they cover a total of 78 environmental types. We grouped these environmental types into 11 categories: air, indoor surface, sediment, freshwater, marine, non-marine saline, biofilm, soil, plant-associated, animal-associated and human-associated (Supplementary table 1).

### Estimation of the average *rrn* copy number (ACN) of a community

The 16S rRNA gene copy number of each OTU in the SILVA and the GreenGene reference databases was estimated using the phylogenetic ancestral state reconstruction method described in Kembel *et al* (Kembel et al., 2012) with an updated reference tree of 1,197 bacterial species whose genomes were sequenced and 16S rRNA gene copy numbers were known. The ACN of the bacterial community in a sample was weighted by the relative cell abundance of OTUs, as shown in Equation (1),

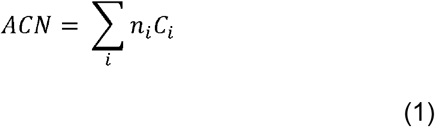

where *n*_*i*_ and *C*_*i*_ are the relative cell abundance and 16S rRNA gene copy number, respectively. The relative cell abundances *n*_*i*_ was in turn computed by Equation (2) proposed in Kembel *et al* (Kembel et al., 2012).

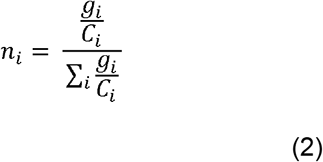

where g_i_ is the relative gene abundance of an OTU estimated by the read number.

### Estimation of the distribution of *rrn* copy number in communities of an environment type

The relative cell abundance of bacteria with a certain *rrn* copy number was calculated by summing the relative cell abundance of all species in a community with that *rrn* copy number (rounded to the nearest integer). To generate the distribution plot for an environmental type (Figure 2C), we repeated the analysis for all samples (communities) of that environmental type and calculated the average relative cell abundance of bacterial species with a certain *rrn* copy number.

## Competing interests

The authors declare no competing financial interests.

## Supplementary Data

**Supplementary Figure 1.**
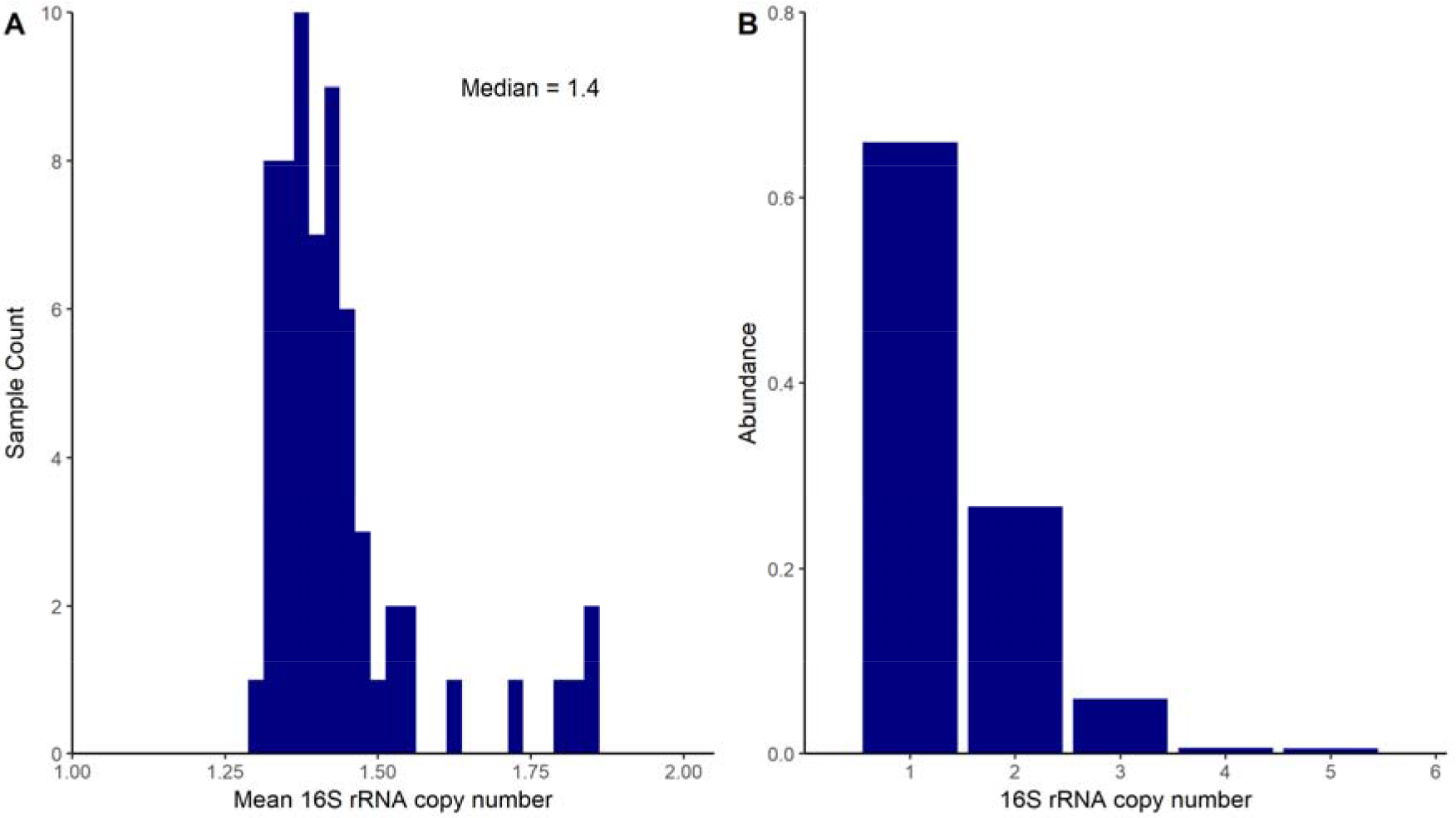
(A) The average copy number of *rrn* copy of the 63 bacterial communities from the surface layer in TARA. (B) The distribution of *rrn* copy number in the 63 bacterial communities. The abundances of species with more than 6 *rrn* copies are negligible and not shown.

Supplementary Table 1. Environmental types covered by samples from WGS metagenomics in the EBI Metagenomics (A) and 16S rRNA amplicon sequencing in the EBI Metagenomics and the Earth Microbiome Project (B), and their statistics.

Supplementary Table 2. The *rrn* copy number of 116 strains of oligotrophic or copiotrophic bacteria whose complete genome sequences were available.

